# Lymphopenia drives T cell exhaustion in immunodeficient STING gain-of-function mice

**DOI:** 10.1101/2024.11.01.621470

**Authors:** Damien Freytag, Stéphane Giorgiutti, Nadège Wadier, Sabine Depauw, Grégoire Hopsomer, Philippe Mertz, Fabrice Augé, Raphaël Carapito, Isabelle Couillin, Anne-Sophie Korganow, Francesca Pala, Marita Bosticardo, Luigi D. Notarangelo, Frédéric Rieux-Laucat, Nicolas Riteau, Peggy Kirstetter, Pauline Soulas-Sprauel

## Abstract

STING gain-of-function (GOF) mutations are associated with the severe autoinflammatory disease designated STING Associated Vasculopathy with onset in Infancy (SAVI). Mice with the STING GOF V154M mutation develop profound T cell lymphopenia, partly due to a blockage of T cell development in the thymus. To better characterize the mechanisms of peripheral T cell dysfunctions, we conducted a transcriptomic and phenotypic analysis on sorted splenic CD4+ and CD8+ mature T cells from STING GOF V154M mice. We identify a T cell exhaustion phenotype that manifests at a terminal stage, acquired early in life but only after reaching the peripheral environment. This phenotype is independent of type I interferons and does not rely on intrinsic STING activation in either T cells or stromal cells. Mechanistically, the limited number of mature T cells that reach the periphery appear to be quickly impacted by the lymphopenic environment, experiencing heightened stimulation of the IL-7 receptor and TCR pathways, including the NFAT pathway, a key factor in T cell exhaustion. By performing transplantation experiments with STING GOF long term-hematopoietic stem cells (LT-HSCs) along with supportive wild-type bone marrow (BM) cells, we were able to prevent the T cell exhaustion of STING GOF T cells in the resulting non-lymphopenic context, demonstrating that lymphopenia is a major driver of T cell exhaustion in STING GOF mice. T cell exhaustion, although less severe, was also observed in lymphopenic mice carrying *Rag1* hypomorphic mutations. In conclusion, our results, which highlight T cell exhaustion induced by lymphopenia, could have important implications for the management of patients with severe immune deficiencies.

**Highlights:**

- We describe a phenotype of T cell exhaustion in STING GOF V154M mice, which is acquired early in life and in the periphery.
- STING GOF-associated T cell exhaustion is independent of type I IFNs, and STING GOF/activation in T cells or in stromal cells is not sufficient for T cell exhaustion.
- Lymphopenia is a major driver of T cell exhaustion in STING GOF mice, and increased antigenic/IL-7 stimulation of T cells in the lymphopenic context of STING GOF mice could be implicated in the induction of T cell exhaustion.

## Introduction

Stimulator of Interferon Genes (STING) belongs to cytosolic double-stranded-DNA-sensing pathways in innate immunity and has emerged as a central player in antiviral, antibacterial, and anti-tumor immunity (1). When activated by cyclic dinucleotides produced by cyclic GMP-AMP synthase (cGAS), STING translocates from endoplasmic reticulum (ER) to ER-Golgi intermediate compartment (ERGIC)/Golgi where it activates TANK-binding kinase 1 (TBK1), which then phosphorylates interferon regulatory factor 3 (IRF3) and induces the expression of type I interferons (IFNs). In turn, type I IFNs activate the type I IFNs receptor (IFNAR), inducing the expression of a set of interferon-stimulated genes (ISG) and triggering inflammation. The NF-kB pathway is also engaged following STING activation, participating in the inflammatory response (2).

Heterozygous gain-of-function (GOF) mutations in STING-coding gene *STING1* lead to constitutive activation of the protein and have been described in patients with an autoinflammatory disease designated STING Associated Vasculopathy with onset in Infancy (SAVI) (3, 4), classified as a type I interferonopathy, with increased ISGs expression (5). This disease is associated with damages to the skin small blood vessels and an interstitial lung disease at the origin of morbidity and mortality (6–8). To study the role of STING in the pathophysiology of the disease, our team (9) and others (10–16) developed STING GOF mouse models including mice carrying the heterozygous V154M mutation (9, 10), corresponding to the most common human V155M mutation (about 60%) in SAVI patients. STING GOF V154M mouse models develop type I IFN-independent auto-inflammatory manifestations, notably lung disease (10, 17), and also a high decrease of T cell counts in a context of severe combined immunodeficiency (SCID) (9, 10), reminiscent of T cell lymphopenia cases among SAVI patients. T cell lymphopenia was also described in STING GOF N153S models (10, 11). In STING GOF N153S and V154M mice, the T cell lymphopenia is partly explained by an impairment of T cell development at double negative (DN)1 and DN2 stages (9, 11).

In addition, we showed that the rare mature T cells found in the periphery exhibited both defective proliferation and increased apoptosis, which maintain the lymphopenic state (9). These defects are at least partly STING-intrinsic as its anti-proliferative effect was confirmed in human T cells overexpressing the STING GOF V155M mutant (18). In addition, a pro-apoptotic effect of STING in lymphocytes was also reported in other models, and involves TBK1 (19), endoplasmic reticulum (ER) stress (20) or p53-dependent pathways (21). However, treatment of STING N153S mice with a pan-caspase inhibitor or ER stress blocker only partially reverse T cell lymphopenia (20). Thus, other mechanisms must explain the functional defects of peripheral T cells and maintenance of lymphopenia in STING GOF mice.

To better characterize these mechanisms, we perform in the present study a transcriptomic analysis on sorted T cells from STING GOF mice compared to their wild-type (WT) littermate controls. An enrichment of transcriptomic signatures associated with T cell exhaustion is revealed and confirmed by the overexpression of key transcription factors and several surface inhibitory receptors (like PD-1, TIGIT, TIM-3 and LAG-3). We further demonstrate that T cell exhaustion is associated with enhanced TCR (including NFAT signaling) and IL-7R signaling in the lymphopenic environment. Importantly, we do not observe any sign of T cell exhaustion in STING GOF T cells developing in non-lymphopenic mice. Finally, a similar T cell exhaustion phenotype (albeit less severe than in STING GOF mice) is observed in other lymphopenic conditions, as shown in mice carrying hypomorphic mutations of *Rag1* gene (22). Together, these observations demonstrate that T cell exhaustion phenotype in STING V154M mice is mostly triggered by lymphopenia and likely reinforce this lymphopenia by a self-maintaining loop. While T cell exhaustion has primarily been described as a consequence of chronic T cell stimulation in the context of persistent infections and cancers (23, 24), our study reveals that lymphopenia should also be considered as a context that can trigger T cell exhaustion, echoing recent findings of T cell exhaustion in patients with severe combined immunodeficiencies (SCID) (25, 26).

## Results

### CD4^+^ and CD8^+^ T cells from STING GOF mice display an exhausted phenotype, acquired in the periphery in a type I IFN independent manner

Mice carrying the heterozygous STING V154M mutation are characterized by a SCID phenotype associated with marked T cell dysfunctions, including impaired proliferation, and enhanced apoptosis (9). To decipher the mechanisms underlying this phenotype, we performed a transcriptomic analysis by RNA-seq on sorted splenic CD4^+^ and CD8^+^ T cells from STING GOF mice compared to their WT littermate controls. This analysis revealed a transcriptomic signature of T cell exhaustion in CD4^+^ and CD8^+^ T cells in the STING GOF group, as shown by a negative enrichment of the transcripts upregulated in naive versus exhausted T cells, as well as a positive enrichment of the genes downregulated in naive versus exhausted T cells (GSE9650 (27)) (**Fig. 1A**). T cell exhaustion is an altered differentiation state of chronically activated T cells, notably caused by persistent antigenic stimulation and sustained inflammation, and is commonly occurring in chronic viral infections or cancers (23, 24). Exhausted T cells co-express inhibitory receptors (like PD-1, TIGIT, TIM-3 and LAG-3) in a sustained manner (24). Furthermore, exhausted T cells progressively lose their proliferative and effector functions, which is reminiscent of the functional defects of mature STING GOF T cells (9, 10).

**Figure 1.**
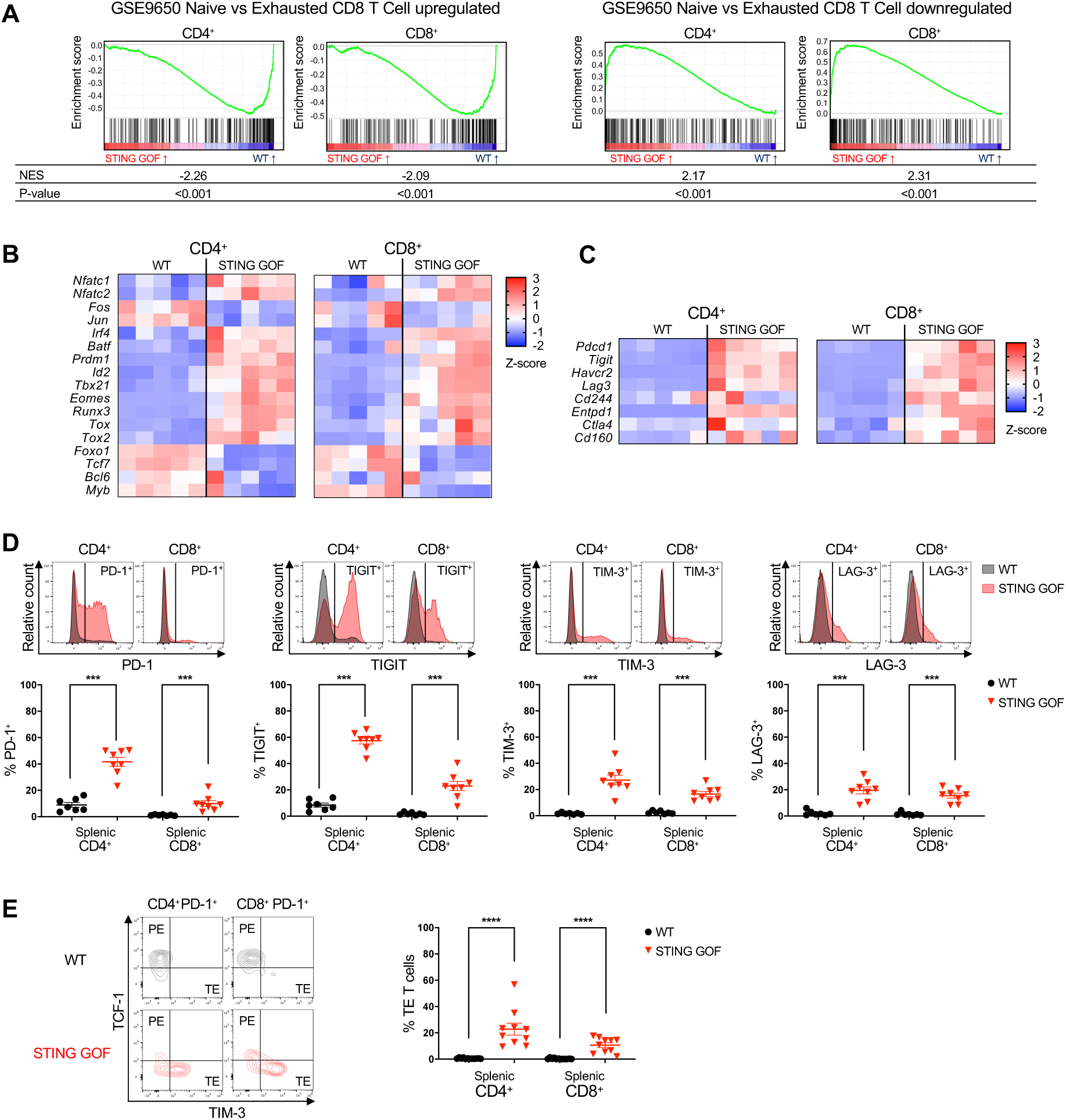
Splenic CD4+ and CD8+ T cells from STING GOF mice display an exhausted phenotype. **(A-C)** Analysis of RNA-seq performed on sorted splenic CD4^+^ or CD8^+^ T cells from STING GOF mice (n=5) and their WT littermate controls (n=5). **(A)** Gene Set Enrichment Analysis (GSEA) of the signatures of genes upregulated or downregulated in naive versus exhausted CD8^+^ T cells (GSE9650) among genes deregulated in STING GOF versus WT CD4^+^ or CD8^+^ T cells. Enrichment plots, normalized enrichment score (NES) and nominal p-value (p-value) are shown for each analysis. **(B)** Heatmap representing T cell exhaustion/dysregulated transcription factors mRNA expression (Z-score) by CD4^+^ or CD8^+^ T cells in STING GOF mice compared to WT mice. **(C)** Heatmap representing inhibitory receptors mRNA expression (Z-score) by CD4^+^ or CD8^+^ T cells in STING GOF and WT mice. **(D-E)** Immunophenotyping of splenic T cells from STING GOF mice and their WT littermate controls by flow cytometry. **(D)** Proportion of PD-1-, TIGIT-, TIM-3- and LAG-3-expressing cells among CD4^+^ or CD8^+^ T cells. Representative histograms are shown. **(E)** Proportion of terminally exhausted (TE) T cells among CD4^+^ or CD8^+^ T cells. Representative contour plots are shown. Each data point corresponds to one mouse; mean ± SEM are shown per population for seven to twelve mice from multiple independent experiments. Statistical significances are calculated with two-tailed Mann-Whitney test: ***, P < 0.001; ****, P < 0.0001.

Analysis of gene expression for key transcription factors associated with T cell exhaustion in STING GOF T cells revealed overexpression of *Nfatc1 and Nfatc2* compared to WT cells, without concurrent induction of the AP-1 cofactors *Fos* and *Jun* (**Fig. 1B**). This pattern is indicative of a preferential activation of the transcriptional program of exhaustion (28). We also observed increased expression of T cell exhaustion-associated transcription factors, like EOMES (*Eomes*) and T-BET (*Tbx21*), as well as TOX (*Tox*), the latter being more involved in the most advanced stages of T cell exhaustion program (**Fig. 1B**). Moreover, transcripts of multiple genes encoding for inhibitory receptors (*Pdcd1*, *Tigit*, *Havcr2*, *Lag3*, *Cd244*, *Entpd1*, *Ctla4* and *Cd160*) were also overexpressed (**Fig. 1C**). Accordingly, the percentages of splenic CD4^+^ and CD8^+^ T cells positive for PD-1, TIGIT, TIM-3 or LAG-3 expression were significantly increased in STING GOF mice compared to their WT controls (**Fig. 1D**). Then, we analyzed the expression of additional markers to define more precisely the exhaustion stage of T cells in STING GOF mice. As *Tcf7* transcript was downregulated in our RNA-seq data (**Fig. 1B**), we hypothesized that STING GOF T cells could be terminally exhausted. We analyzed co-expression of the inhibitory receptors PD-1 and TIM-3, as well as the transcription factor TCF-1 (encoded by *Tcf7* gene), to identify progenitor exhausted (PE) T cells (PD-1^+^TCF-1^+^TIM-3^-^) and terminally exhausted (TE) T cells (PD-1^+^TCF-1^-^TIM-3^+^) (29). An increased proportion of TE T cells was observed among CD4^+^ and CD8^+^ T cells from STING GOF mice as compared to control mice. Interestingly, the STING GOF T cell exhaustion phenotype appeared to be more profound in CD4^+^ than in CD8^+^ T cells (**Fig. 1E**). Together, our findings revealed an unexpected phenotype of T cell exhaustion in STING GOF mice, which reaches the stage of terminal exhaustion, and might contribute to the T cell dysfunctions described in these mice.

Given that STING V154M mice develop a lung disease partly mediated by T cells (17), we also investigated potential T cell exhaustion in the lungs. Interestingly, as in the spleen, the proportion of CD4^+^ and CD8^+^ T cells infiltrating the lungs that expressed PD-1 (data available for CD4^+^ T cells), TIGIT, TIM-3 or LAG-3 was significantly higher in STING GOF mice compared to WT controls (**Fig. S1A**).

To determine if this exhaustion phenotype has a central origin, we examined whether the single positive (SP) T cells, the most terminally differentiated T cell progenitors in the thymus, were already exhibiting the phenotype. We observed only minimal increase of cells expressing inhibitory receptors, for both thymic SP CD4^+^ and CD8^+^ progenitors (**Fig. S1B**). In contrast, we demonstrated that splenic T cells from very young (2-week-old) mice, which represent cells that have just left the thymus for the periphery, were already displaying the exhaustion phenotype of adult mice (**Fig. S1C**). Altogether, these results demonstrate that the T cell exhaustion phenotype in STING GOF mice is acquired very early after birth, but only once T cells have reached the periphery.

Since STING drives type I IFNs, which have recently shown to regulate coinhibitory receptors expression on T cells (30), we then explored the role of these cytokines in T cell exhaustion in STING GOF mice. Overall, the T cell exhaustion phenotype was not reversed by IFNAR deficiency in STING GOF mice (**Fig. S2**).

### Ca^2+^-NFAT pathway is activated by STING, but is not sufficient to induce T cell exhaustion

We then searched for signals that could induce the exhaustion phenotype in STING GOF T cells. Considering its crucial role in T cell exhaustion (31) and its upregulation in our RNA-seq data (**Fig. 1B**), we first focused on NFAT signaling pathway.

Transcriptomic analysis confirmed a significant enrichment of NFAT signature from the Pathway Interaction Database (PID) in STING GOF T cells, compared to control T cells (**Fig. S3A**). Upregulation of the total (nuclear and cytosolic) NFATc1 transcription factor expression was confirmed by flow cytometry (**Fig. S3B**). NFAT activation relies on its dephosphorylation by the calcium (Ca^2+^)-calmodulin-calcineurin pathway, and Ca^2+^ homeostasis is known to be disrupted in STING GOF T cells (20). Using Fura Red™-mediated ratiometric measurement to study basal Ca^2+^ levels, we demonstrated that basal Ca^2+^ levels were higher in CD4^+^ and CD8^+^ T cells from STING GOF mice compared to their WT controls, suggesting that Ca^2+^-NFAT signaling pathway in STING GOF T cells participates to the induction of their exhaustion (**Fig. S3C**).

We next wondered whether the activation of the Ca^2+^-NFAT signaling pathway could be due to the STING activation *per se*. Therefore, we treated WT splenocytes with STING agonist 5,6-dimethylxanthenone-4-acetic acid (DMXAA) *in vitro*. After 4 hours, DMXAA-treated CD4^+^ and CD8^+^ T cells presented an increase of their basal Ca^2+^ levels, similar to those observed in untreated STING GOF T cells (**Fig. 2A**). After 24 hours, DMXAA-treated cells also showed upregulation of NFATc1 to the same extent as untreated STING GOF T cells (**Fig. 2B**). However, STING activation *in vitro* was not sufficient to induce an increase of key inhibitory receptors PD-1 and TIM-3 (**Fig. 2C**).

**Figure 2.**
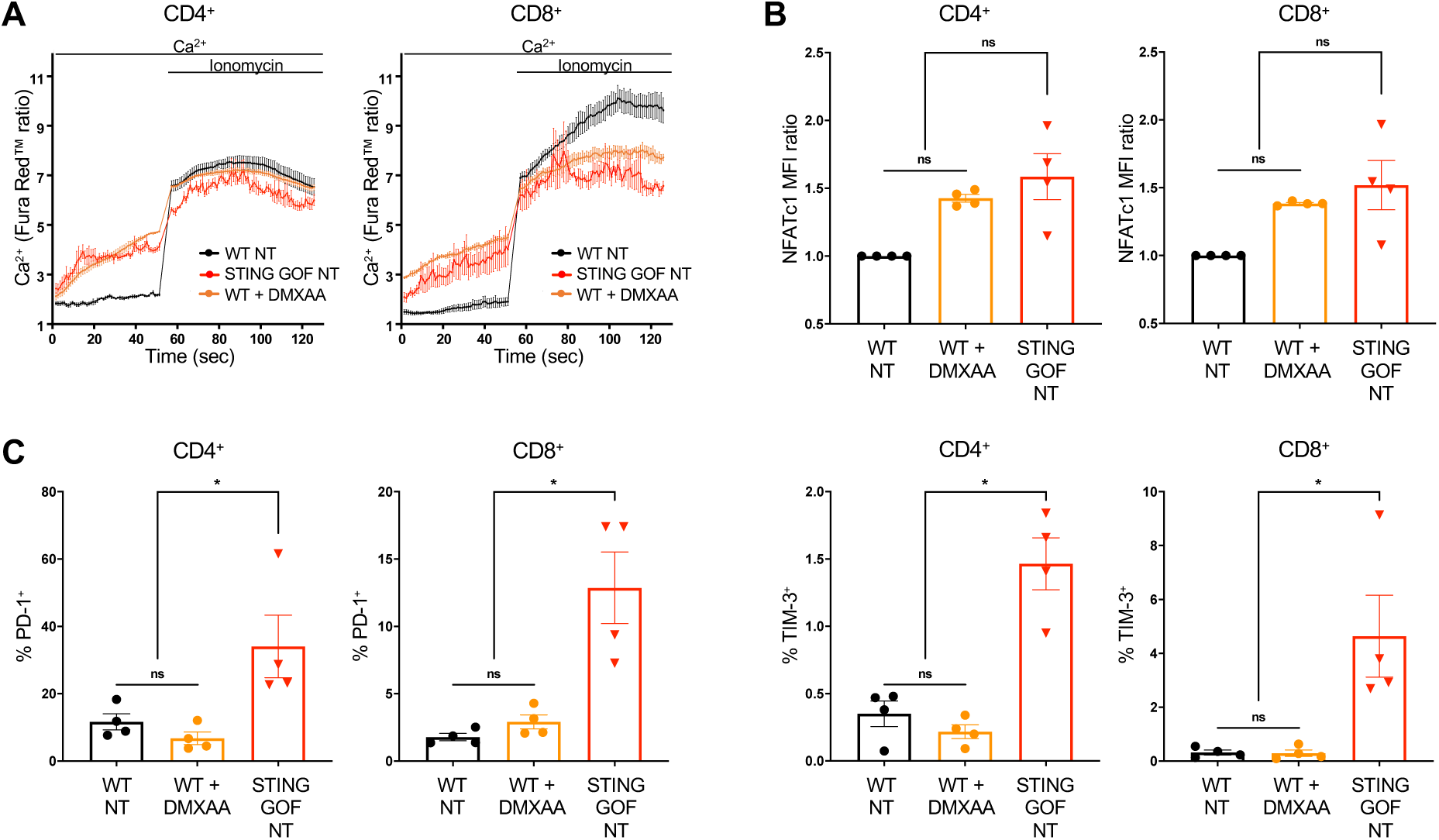
Ca^2+^-NFAT pathway is activated by STING, but is not sufficient to induce T cell exhaustion. **(A-C)** Splenic WT splenocytes were treated *in vitro* with DMXAA (10 µg/mL) and compared to untreated (NT) WT and STING GOF splenocytes. After 4 hours (calcium levels measurement) or 24 hours (immunophenotyping), cells were analyzed by flow cytometry. **(A)** Relative cytosolic Ca^2+^ levels monitored by Fura Red™ ratio in CD4^+^ or CD8^+^ T cells Splenocytes were treated, recorded, and plotted as in Fig. S3C. Each data point represents the mean of three independent experiments, and their error bars the SEM. **(B)** Ratio of total NFATc1 MFI in CD4^+^ or CD8^+^ T cells. Ratio was normalized on the corresponding WT untreated control. **(C)** Proportion of PD-1- and TIM-3-expressing cells among CD4^+^ or CD8^+^ T cells. Each data point corresponds to one mouse; mean ± SEM are shown per population for four mice from four independent experiments. For NFATc1 MFI ratio, statistical significances are calculated with Wilcoxon signed-rank test with a hypothetical value of 1: ns, P > 0.05. For PD-1^+^ and TIM-3^+^ proportions, statistical significances are calculated with two-tailed Mann-Whitney test: ns, P > 0.05; *, P < 0.05.

In conclusion, even if it can contribute to the Ca^2+^-NFAT activation, STING activation in T cells is not sufficient to induce their exhaustion, reinforcing the importance of an environmental factor that was already suggested comparing thymic versus splenic exhaustion profile from 2-weeks old STING GOF mice (**Fig. S1C**).

### STING GOF T cells displayed features of TCR and IL-7R engagement mediated by the lymphopenic environment

We then hypothesized that NFAT activation in STING GOF could be the consequence of TCR signaling secondary to antigenic stimulations (31). Indeed, beyond NFAT overexpression, transcriptomic analysis confirmed a significant enrichment of a broader TCR signaling pathway signature from the Wikipathways (WP) in STING GOF T cells compared to control T cells (**Fig. 3A**). In addition, we highlighted a significant downregulation of CD3 expression (consistent with TCR stimulation) on the surface of STING GOF T cells compared to control T cells (**Fig. 3B**).

**Figure 3.**
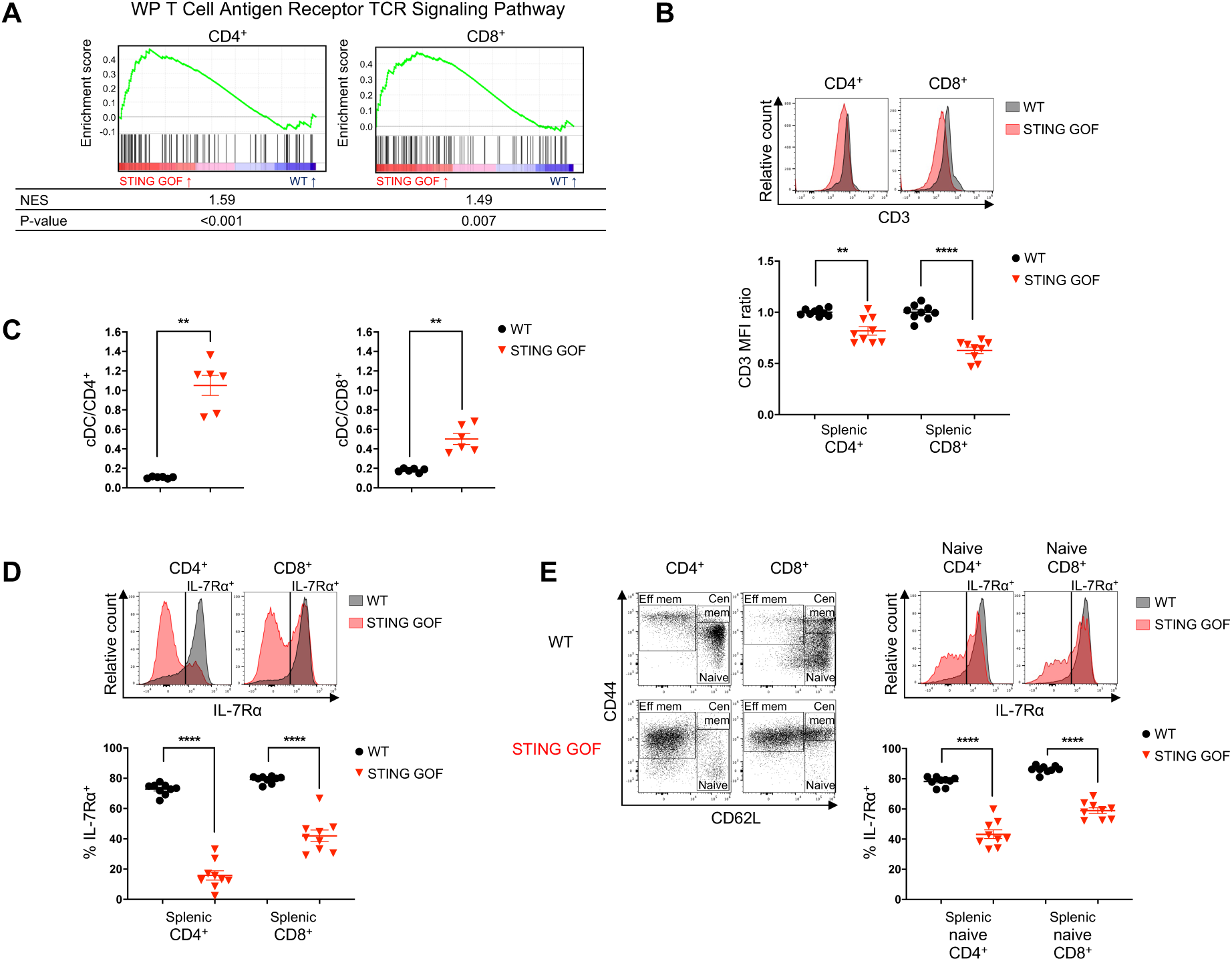
STING GOF T cells displayed features of TCR and IL-7R engagement mediated by the lymphopenic environment. **(A)** GSEA of the TCR pathway signature (WP) among genes deregulated in STING GOF versus WT CD4^+^ or CD8^+^ T cells. Enrichment plots, normalized enrichment score (NES) and nominal p-value (p-value) are shown for each analysis. **(B)** Ratio of CD3 mean fluorescence intensity (MFI) on splenic CD4^+^ or CD8^+^ T cells from STING GOF mice and their WT littermate controls. Ratio was normalized on the mean of WT controls of each analysis. Representative histograms are shown. **(C)** Ratio of cDCs absolute number per CD4^+^ or CD8^+^ T cells numbers from the spleen of 4-week-old mice. **(D-E)** Proportion of IL-7Rα-expressing cells among splenic total **(D)** and naive (CD44^-^CD62L^+^) **(E)** CD4^+^ or CD8^+^ T cells from STING GOF mice and their WT littermate controls. Gating strategy of naive T cells and representative histograms are shown. Each data point corresponds to one mouse; mean ± SEM are shown per population for six to nine mice from multiple independent experiments. Statistical significances are calculated with two-tailed Mann-Whitney test: **, P < 0.01; ****, P < 0.0001.

This antigenic stimulation was not secondary to an acute or chronic infection status, as STING GOF mice were bred and held under specific opportunist pathogen free (SOPF) environment, and was not associated with cancer as no tumors were detected in these mice. Thus, we hypothesized that increased TCR stimulation by conventional dendritic cells (cDCs) could be responsible for this phenotype, similar to what is observed during homeostatic proliferation of T cells in lymphopenic environments, promoting the reconstitution of the lymphoid compartment in the periphery (32). Interestingly, we showed a much higher number of cDCs per CD4^+^ T cell, and, to a lesser extent per CD8^+^ T cell, in STING GOF mice compared to their WT controls, which could enhance antigen-mediated TCR activation (**Fig. 3C**).

Besides antigenic stimulation, IL-7 could also affect T cell fate in STING GOF T-cell lymphopenic environment, since IL-7R engagement promote T cell homeostatic proliferation (32). A marked down-regulation of the alpha chain of the receptor (IL-7Rα), reminiscent of IL-7R engagement, was observed for both CD4^+^ and CD8^+^ T cells (**Fig. 3D**). IL-7Rα expression was already significantly downregulated in CD4^+^ and CD8^+^ naive T cells from STING GOF mice (**Fig. 3E**), indicating that T-cell lymphopenia impacts T cell fate at early stage in the periphery of STING GOF mice.

Taken together, our results suggest that the lymphopenic environment impacts T cells in STING GOF mice through TCR and IL-7R engagement, which are known to govern lymphopenia-induced homeostatic proliferation.

### T cell exhaustion phenotype of STING GOF T cell is reversed in a non-lymphopenic environment

We next wondered whether STING GOF T cells would display an exhausted profile in a non-lymphopenic context. To this purpose, we took advantage of bone marrow transplantation experiments. We sorted long term-hematopoietic stem cells (LT-HSCs) from bone marrow

(BM) of donor STING GOF or WT littermate control mice (CD45.2^+^, donor) and co-transplanted them along with supportive WT BM cells (CD45.1^+^, support) into irradiated WT recipient mice (CD45.1^+^CD45.2^+^, host) (**Fig. 4A**). First, we confirmed the hematopoietic intrinsic effect of the STING GOF mutation on splenic T cell reconstitution (10), as very few T cells were derived from the STING GOF LT-HSCs (red dots), while reconstitution was effective from WT LT-HSCs (black dots). However, the lack of T cell reconstitution from the STING GOF LT-HSCs was compensated by transplantation of supportive WT BM-derived T cells (grey dots) (**Fig 4B**). Consequently, absolute numbers of total (including both CD45.1^+^ and CD45.2^+^) CD4^+^ and CD8^+^ T cells were similar in STING GOF (STING GOF➔WT) and WT LT-HSCs (WT➔WT) transplantation settings (**Fig 4C**). Interestingly, STING GOF LT-HSCs-derived CD4^+^ and CD8^+^ T cells did not overexpress the inhibitory receptors PD-1, TIGIT, TIM-3 or LAG-3 when compared to WT LT-HSCs-derived T cells, or to WT supportive BM-derived T cells (**Fig 4D**). Consistent with this, there were no terminally exhausted (TE) T cells among STING GOF LT-HSC-derived CD4^+^ and CD8^+^ T cells (**Fig 4E**).

**Figure 4.**
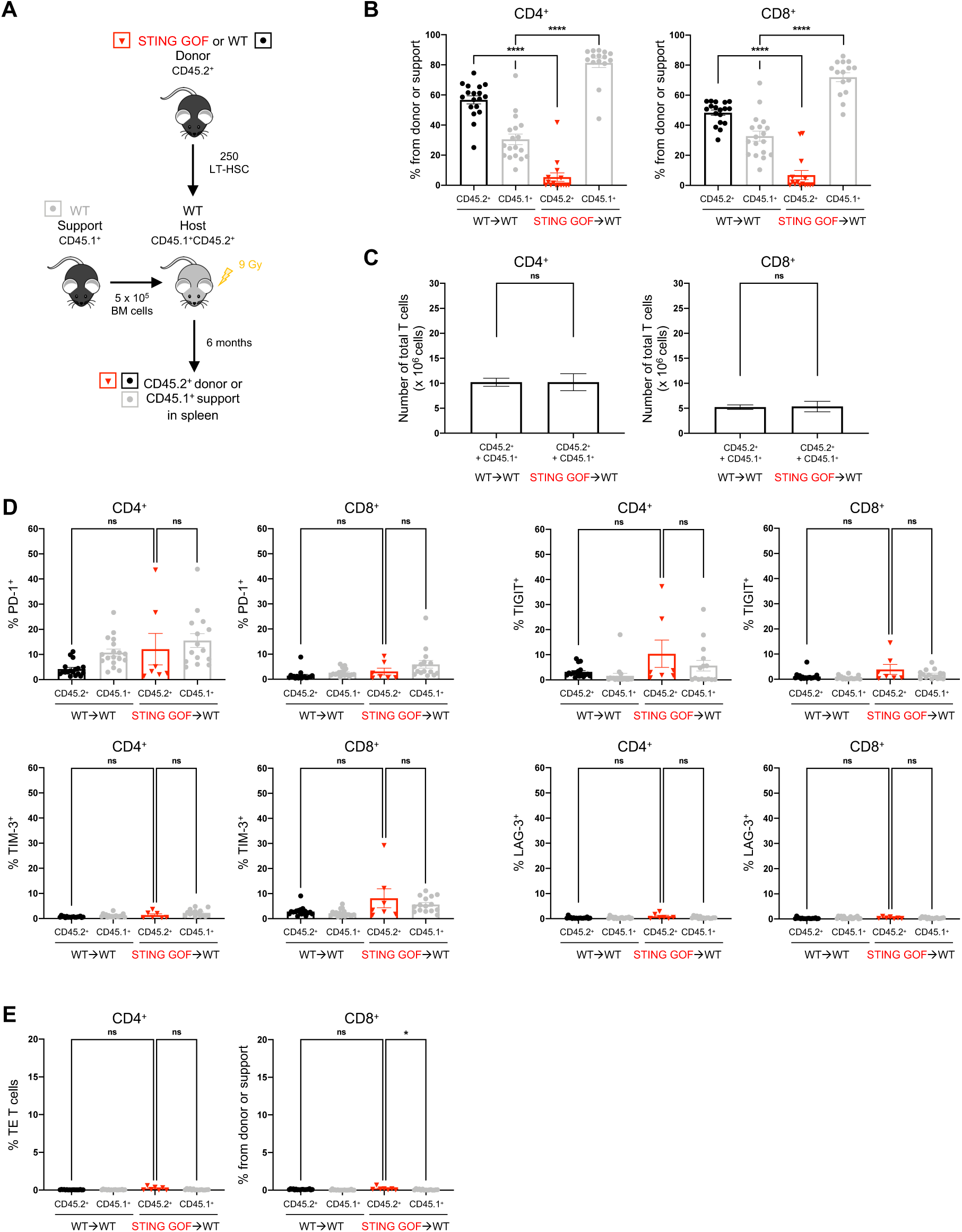
T cell exhaustion phenotype of STING GOF T cell is reversed in a non-lymphopenic environment. **(A-E)** LT-HSCs from STING GOF mice and their WT littermate controls (CD45.2^+^) were transplanted with WT BM supportive cells (CD45.1^+^) in WT irradiated hosts to generate STING GOF➔WT and WT➔WT mice. 6 months later, spleens were assessed for T cell reconstitution and immunophenotyping. **(A)** Strategy of LT-HSCs transplantations with supportive BM cells. **(B)** Proportion of cells derived from donor (CD45.2^+^) LT-HSCs or WT BM supportive cells (CD45.1^+^) among splenic CD4^+^ or CD8^+^ T cells from STING GOF➔WT and WT➔WT mice. **(C)** Absolute numbers of total (CD45.2^+^ and CD45.1^+^) splenic CD4^+^ or CD8^+^ T cells from STING GOF➔WT and WT➔WT mice. **(D)** Proportion of PD-1-, TIGIT-, TIM-3- and LAG-3-expressing cells among splenic CD4^+^ or CD8^+^ T cells **(E)** Proportion of terminally exhausted (TE) T cells among splenic CD4^+^ or CD8^+^ T cells. Each data point corresponds to one mouse; mean ± SEM are shown per population for seven to eighteen mice from multiple independent experiments. Statistical significances are calculated with two-tailed Mann-Whitney test: ns, P > 0.05; *, P < 0.05; ****, P < 0.0001.

Nevertheless, because of the WT background of recipient mice in LT-HSC transplantations, radio-resistant stroma was composed of WT cells (**Fig 4A**), which is not the case in STING GOF mice. In order to determine the potential impact of the STING GOF stroma on the T cell exhaustion phenotype, we performed transplantation experiments in which total BM cells from WT donor mice (CD45.1^+^, donor) were transplanted into STING GOF or WT irradiated recipient mice (CD45.2^+^, host) (**Fig. S4A**). As expected, a good splenic T-cell reconstitution were observed upon transplantation of WT donor BM cells into STING GOF irradiated host (**Fig. S4B**). In addition, the absolute numbers of CD4^+^ and CD8^+^ T cells were comparable in the two groups (WT➔WT, and WT➔STING GOF) of mice (**Fig. S4C**). Moreover, the proportion of CD4^+^ and CD8^+^ T cells expressing the inhibitory receptors PD-1, TIGIT, TIM-3 or LAG-3 was similar following transplantation of WT donor BM cells into irradiated STING GOF mice or WT recipients (**Fig. S4D**), and the proportion of TE T cells was extremely low in either group of mice (**Fig. S4E**).

Together, these data indicate that STING GOF stroma does not trigger T cell exhaustion phenotype and rather identify T cell lymphopenia as the main mechanism driving T cell exhaustion in STING GOF mice.

### Mice carrying Rag1 hypomorphic mutations also display lymphopenia-associated T cell exhaustion

In order to determine if lymphopenia-mediated T cell exhaustion in STING GOF mice is specific to the STING GOF mutation, we investigated whether another lymphopenic mouse model might also show T cell exhaustion. To this purpose, we studied two mouse models carrying hypomorphic *Rag1* mutations (*Rag1*^R972Q/R972Q^ and *Rag1*^R972W/R972W^ mice, thereafter referred to as RAG1 R972Q and RAG1 R972W mice), also presenting T cell lymphopenia of central origin, based on early block in T cell development (22) as in STING GOF mice. Indeed, RAG1 R972Q and R972W mutant mice similar levels of CD4^+^ and CD8^+^ T cell lymphopenia in spleen, compared to STING GOF mice (**Fig. S5A**).

Interestingly, a significant proportion of CD4^+^ and CD8^+^ T cells from RAG1 R972Q and R972W mice was expressing inhibitory receptors compared to their WT controls, albeit not to the same extent as observed in STING GOF T mice for the majority of inhibitory receptors studied (**Fig. 5**). TE T cells were also increased among CD4^+^ and CD8^+^ T cells for both RAG1 mutants compared to their WT controls, without reaching the same proportion as in STING GOF mice (**Fig. S5B**). In conclusion, lymphopenia seems to be sufficient to induce T cell exhaustion. However, as T cell exhaustion appeared to be more severe in STING GOF mice compared to RAG1 hypomorphic mutant mice, additional factors beyond central T cell lymphopenia must contribute to the higher degree of T cell exhaustion observed in STING GOF mice.

**Figure 5.**
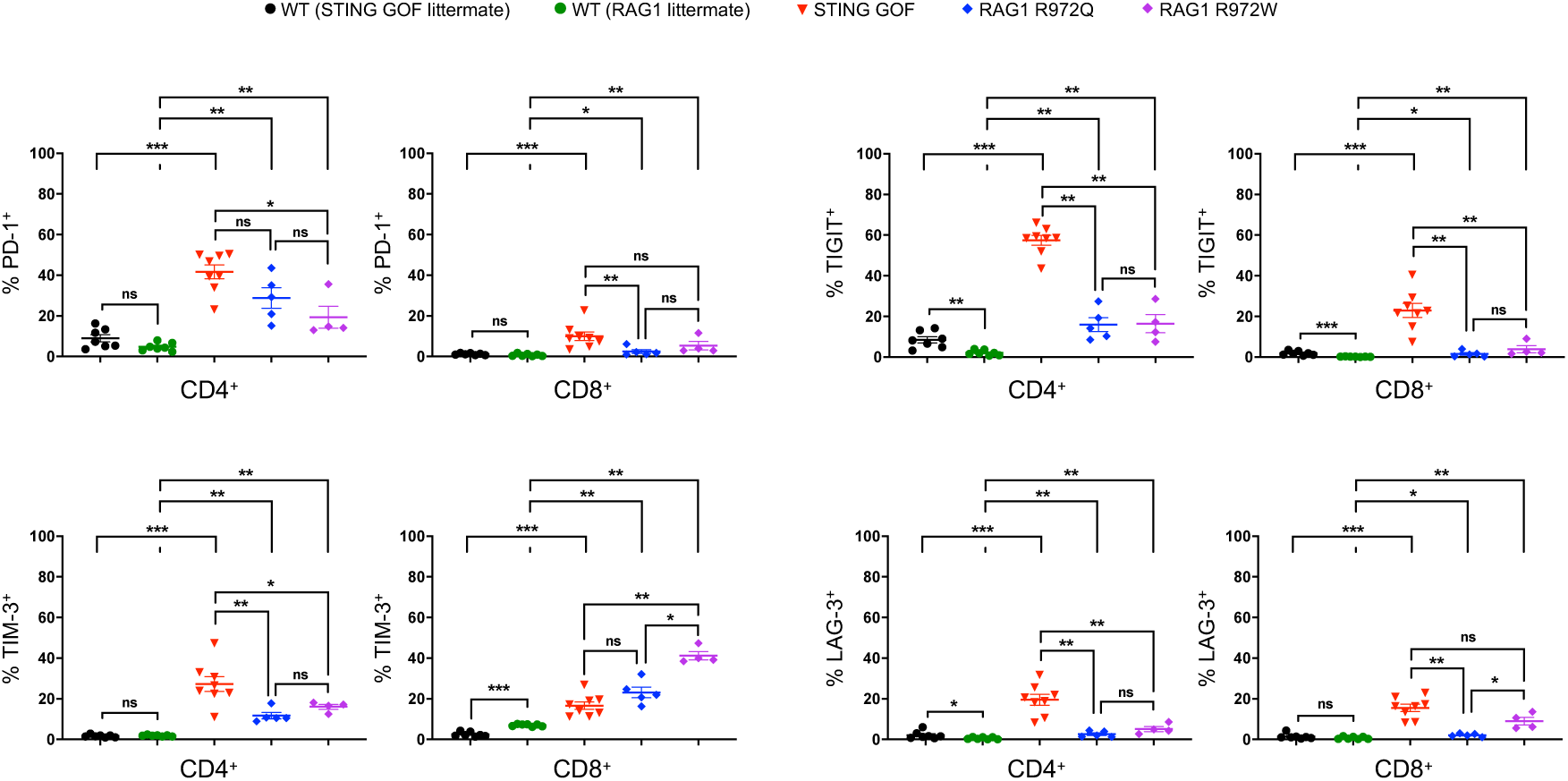
Mice carrying Rag1 hypomorphic mutations also display lymphopenia-associated T cell exhaustion. Proportion of PD-1-, TIGIT-, TIM-3- and LAG-3-expressing cells among splenic CD4^+^ or CD8^+^ T cells from hypomorphic RAG1 R972Q and R972W mice and their WT littermate controls by flow cytometry. Data were compared with previous results obtained from STING GOF mice and their WT littermate controls. Each point corresponds to one mouse; mean ± SEM are shown per population for four to twelve mice from multiple independent experiments. Statistical significances are calculated with two-tailed Mann-Whitney test: ns, P > 0.05; *, P < 0.05; **, P < 0.01; ***, P < 0.001.

## Discussion

Our data revealed an unexpected T cell exhaustion phenotype in STING GOF mice. Usually associated with chronic T cell stimulation (as in the case of chronic viral infections and cancers), this altered state of T cell differentiation turned out to be of particular interest since it could account for some of the functional defects of mature STING GOF T cells, especially their lack of proliferation in response to TCR activation (9). Moreover, we show here that T cell exhaustion in STING GOF mice is type I IFN-independent, as was the T cell proliferation defect (9). Overall, T cell exhaustion could contribute to T cell proliferation defects and apoptosis, in addition to intrinsic effects of the STING protein in mature T cells (18–21).

T cell exhaustion is also associated with a progressive loss of effector properties. In particular, T cell exhaustion may limit IFN-γ production by T cells, which is known to be increased by STING activation (17, 33) and to contribute to the lung disease mortality (17). Therefore, T cell exhaustion could thus be beneficial in STING GOF mice by limiting tissue immunopathology, similarly to the effect of T cell exhaustion in other auto-inflammatory and autoimmune diseases (34). In accordance with this, we also showed an increased proportion of exhausted T cells in lungs of STING GOF mice.

In addition to T cell exhaustion associated with prolonged antigenic stimulations as chronic viral infections or tumoral context (35), we describe here a T cell exhaustion state whose main driver is represented by lymphopenic environment. Like the two other contexts, T cell exhaustion in STING GOF mice appeared to be driven mainly by the environment rather than an intrinsic effect in T cells. Indeed, acute treatment of WT splenocytes with DMXAA *in vitro* had a pre-activating effect on T cells without inducing an exhaustion phenotype. In addition, bone marrow transplantation experiments showed that STING GOF LT-HSCs-derived T cells developing in a predominantly WT and non-lymphopenic environment do not manifest the exhaustion phenotype. Among environmental factors potentially implicated in the T cell exhaustion of STING GOF mice, type I IFNs were shown to be dispensable, despite the potential role of these cytokines suggested in the literature (30, 36, 37). When studying other environmental factors, the STING GOF stroma does not appear to be sufficient to induce T cell exhaustion.

STING GOF T cells display signs of TCR and IL-7R engagement, consistent with lymphopenia-associated signals provided by (auto)antigen and IL-7-mediated stimulation, respectively. While these signals are known to lead to generation of memory-like T cells through homeostatic proliferation (32), T cell exhaustion appeared as a new phenomenon, presumably linked to the persistency of these signals. Of note, the observation that the lymphopenia leads to a higher number of cDCs per CD4^+^ T cells than for CD8^+^ T cells may help explain the more severe T cell exhaustion phenotype observed for CD4^+^ T cells. Finally, the observation that mice carrying hypomorphic *Rag1* mutations are lymphopenic and display obvious signs of T cell exhaustion further supports the notion that lymphopenia may represent an important mechanism leading to T cell exhaustion.

Nevertheless, T cell exhaustion in STING GOF mice appears to be more severe than the one observed in the two RAG1 hypomorphic mutant models studied. This difference does not depend on the severity of the lymphopenia, since the latter was even more profound, especially in the RAG1 R972W mutant. Thus, the STING GOF mutation may synergize with lymphopenia in promoting and aggravating T cell exhaustion. An important contributory mechanism to the phenomenon may be represented by the intrinsic effect of STING on T cell activation, as suggested by the upregulation of Ca^2+^-NFAT pathway upon DMXAA treatment. This NFAT activation had been already highlighted by a previous transcriptomic analysis, also indicating its independence from type I IFN (38). Thus, the STING GOF mutation could place the T cells in a pre-activated state, making them not only more sensitive to endoplasmic reticulum stress and apoptosis (20), but also to exhaustion. Even if we demonstrated that STING GOF radioresistant cells (stroma) do not trigger T cell exhaustion phenotype by themselves, synergistic effects of the STING GOF mutation on the antigen presentation capacity of both non-hematopoietic and hematopoietic cells, especially cDC which are also included in the radioresistant cell population, could reinforce TCR stimulation in the lymphopenic context and thus contribute to exhaustion. The impact of this mutation on major histocompatibility complex (MHC) molecules expression has already been demonstrated in endothelial cells within the lung environment in STING GOF mice (17). Moreover, STING activation by agonists in mice has been associated with the loss of cDC1 (39), a subpopulation that plays a crucial role in maintaining and isolating progenitor exhausted (PE) T cells and preventing their transition into terminally exhausted (TE) T cells (40).

Overall, our data show that the T cell exhaustion phenotype in STING V154M mice is the result of poor central production of T cells and will potentially reinforce this lymphopenia by a self-maintaining loop. These data should be considered in the context of severe T cell lymphopenia as severe combined immunodeficiencies (SCID). Importantly, recent descriptions of SCID patients developing T cell exhaustion have reinforced a lymphopenia-mediated T cell exhaustion mechanism (25, 26). In particular, the analysis of 61 SCID patients receiving hematopoietic cell transplantation (HCT) recently showed that patients with poor T cell reconstitution after HCT display an increased frequency of exhausted T cells, compared to patients with normal T cell counts after HCT, therefore directly linking lymphopenia with T cell exhaustion (25). In addition, T cell depletion treatments were identified as an important factor associated with PD1 expression on T cells in patients receiving HCT (41).

In conclusion, the data and lessons drawn from the STING GOF mice could therefore be of key importance for the follow up of patients receiving HCT on the one hand, but also for a better comprehension of potential T cell exhaustion in lymphopenic murine models on the other hand. Finally, STING GOF mice constitute a good model for the study of T cell exhaustion in non-infectious and non-tumoral contexts.

## Materials and methods

Detailed information on the materials and methods can be found in SI Appendix, Supporting Materials and Methods.

### Mice and cells

Detailed information on mice and cells can be found in SI Appendix, Supporting Materials and Methods.

### Flow cytometry and cell sorting

Splenic, thymic, BM or lung single-cell suspensions were analyzed by flow cytometry according to standards protocol. Cells were stained with the antibodies listed in SI Appendix (Supporting Materials and Methods) in PBS (Gibco) supplemented with 2% v/v FBS (Dutscher). Intracellular of total (nuclear and cytosolic) TCF-1 and NFATc1 stainings were performed with anti-TCF-1 AF647 (S33-966, BD Pharmingen) and anti-NFATc1 PE (7A6, BioLegend) antibodies, after fixation and permeabilization of cells using Foxp3/Transcription Factor Staining Buffer Set (Thermo Fisher Scientific), according to the manufacturer’s instructions. Viability was assessed by Fixable Viability Dye eFluor™ 450 (eBioscience) or DAPI (1 μM/mL Sigma) stainings. Cells were analyzed using an Attune NxT™ flow cytometer (Thermo Fisher Scientific). Data were analyzed using FlowJo software (TreeStar).

### RNA-seq on sorted CD4^+^ and CD8^+^ T cells

Splenocytes from STING GOF mice (n=5) and WT littermate controls (n=5) were stained as described above with anti-CD3e FITC (clone 145–2 C11, BD Biosciences), anti-CD4 AF700 (RM4-5, BD Pharmingen), anti-CD8a PE (53-6 .7, BD Pharmingen) and DAPI (1 μM/mL, Sigma) before positive sorting of live CD3^+^CD4^+^ or CD3^+^CD8^+^ T cells using a BD FACSAria™ Fusion flow cytometer (IGBMC Flow Cytometry Facility, Strasbourg, France). Isolated cells were at least 95% pure. Total RNA was extracted using RNeasy® Plus Micro Kit (Qiagen) according to the manufacturer’s instructions. RNAseq analyses were performed by the Genomax Facility (INSERM U1109, ImmunoRhumatologie Moléculaire, Université de Strasbourg, France). Libraries were prepared from 10 ng RNA using SMARTer Stranded Total RNA-Seq Kit, Pico Input Mammalian (Takara) following the manufacturer’s instructions. Briefly, random primers were used for first-strand synthesis, and ribosomal cDNA was cleaved by ZapR V.2 in the presence of mammalian R-probes V.2. Libraries were pooled and sequenced (paired-end 2×75 bp) on a NextSeq 500 using the NextSeq 500/550 High Output Kit V.2 according to the manufacturer’s instructions (Illumina). For each sample, quality control was carried out and assessed with the next-generation sequencing (NGS) Core Tools FastQC. Reads were aligned against *Mus musculus* mm10 reference genome using TopHat 2 Aligner (42) and gene expression levels were estimated using Cufflinks V.2.1.1 (43). Differential expression analysis was performed with Cuffdiff V.2.1.1 after exclusion of fragments per kilobase million below 9. Gene Set Enrichment Analysis (GSEA) was carried out according to the developer’s recommendations. The STING GOF versus WT FPKMs of all genes detected in the RNA-seq were used for the ranked genes list and compared with the indicated signatures from the Molecular Signatures Database (MSigDB), using GSEA Software (UC San Diego, Broad Institute) (44, 45). Raw RNAseq data have been deposited in the EMBL-EBI ArrayExpress archive (accession no. E-MTAB-13658).

### Cytosolic Ca^2+^ level measurements

For cytosolic Ca^2+^ level measurements, splenic single-cell suspensions were stained as described above with Fixable Viability Dye eFluor™ 450 (eBioscience), anti-CD4 AF700 (RM4-5, BD Pharmingen) and anti-CD8a PE-CD584 (53-6 .7, BD Pharmingen) before being loaded with 2 μM Fura Red™ acetoxymethyl ester (AM) (Thermo Fisher Scientific) in PBS 2% FBS containing 0.02% pluronic F-127 (Thermo Fisher Scientific) for 30 min at 37°C, avoiding light. Loaded cells were washed, resuspended in DMEM (Gibco) containing 1.8 mM Ca^2+^ and incubated for 10 min at 37°C avoiding light, before acquisition using an Attune NxT™ flow cytometer (Thermo Fisher Scientific). After an initial 50-sec acquisition sequence, ionomycin (1 µg/mL, InvivoGen) was added, as positive control, before the acquisition of a second 60-sec sequence. Ca^2+^ levels were determined according to time by ratiometric analyzes between VL4 (405 nm-excitation; 660 nm-emission) and BL3 (488 nm-excitation; 695 nm-emission) channels using FlowJo software (TreeStar).

### Splenocyte culture

*In vitro* splenocytes stimulations were performed in complete RPMI-1640 medium containing L-glutamine (Lonza) supplemented with 10% v/v FBS (Dutscher), 50 mmol/L β-mercaptoethanol (Gibco), 1% penicillin/streptomycin (Gibco), and 10 mmol/L HEPES (Lonza). Splenocytes were stimulated for the indicated times at 37°C, under 5% CO_2_, with the STING agonist 5,6-dimethylxanthenone-4-acetic acid (DMXAA) (10 µg/mL, InvivoGen).

### LT-HSCs transplantations with supportive BM cells

For LT-HSCs sorting, total BM cells were stained for lineage marker expression with biotinylated anti-CD3 (17A2, BioLegend), anti-CD4 (GK1.5, BioLegend), anti-CD8 (53.6.7, eBioscience), anti-CD11b (M1/70, BioLegend), anti-CD19 (1D3, BioLegend), anti-CD45R/B220 (RA3-6B2, BioLegend), anti-CD49b (DX5, BioLegend), anti-Gr-1 (Ly-6G/C) (RB6-8C5, BioLegend), anti-TER-119 (TER-119, BioLegend), before depletion with Dynabeads sheep anti-rat IgG beads (Thermo Fisher Scientific). Lineage-negative cells were stained as described above with Streptavidin AF700 (Invitrogen), anti-CD48 FITC (HM48-1, BioLegend), anti-CD117 (c-Kit) APC (2B8, BioLegend), anti-CD150 (SLAM) PE (TC15-12F12.2, BioLegend), anti-Sca-1 (Ly-6A/E) PE-Cy7 (D7, BioLegend) and DAPI (1 μM/mL, Sigma) before positive sorting of live LT-HSCs (CD48^−^CD150^+^Lin^-^Sca1^+^cKit^+^) using a BD FACSAria™ Fusion flow cytometer (IGBMC Flow Cytometry Facility, Strasbourg, France). Isolated LT-HSCs were at least 98% pure. 250 STING GOF or WT LT-HSCs from 6-weeks old CD45.2^+^ donor mice were i.v. injected with 5 × 10^5^ supporting CD45.1^+^ WT BM cells into 9-Gy lethally irradiated CD45.1^+^CD45.2^+^ WT hosts. The spleens of the mice were harvested 6 months after transplantation and the splenocytes analyzed by flow cytometry as described above.

### Wild-type (WT) total bone marrow cells transplantations

1 × 10^6^ WT total BM cells from 10 to 15-weeks old CD45.1^+^ donor mice were i.v. injected into 9-Gy lethally irradiated CD45.2^+^ STING GOF or WT hosts. Mouse spleens were harvested 6 months after transplantation and the splenocytes analyzed by flow cytometry as described above.

### Statistical analysis

All data are presented as means +/- SEMs. Statistical analyses were performed with Prism 8.2.1 software (GraphPad). Statistical significance was calculated with the non-parametric test of Mann-Whitney test, except for NFATc1 MFI ratio, where statistical significance was calculated with Wilcoxon signed-rank test with a hypothetical value of 1 (see details in each Figure legends).

## Acknowledgements

We thank Fabrice Augé, Christian Galmiche, Delphine Lamon, Manon Lecointe, Fabien Lhericel, Sophie Reibel-Foisset for excellent animal care. We thank Vincent Gies, Frédéric Gros, Aurélien Guffroy, Sophie Jung and Thierry Martin (INSERM UMR - S1109, Strasbourg University, France), Vincent Flacher and Christopher Mueller (CNRS UPR 3572, Institute of Molecular and Cellular Biology (IBMC), Strasbourg, France) for scientific discussions. We thank the Flow Cytometry facility of IGBMC (Illkirch, France) for cell sorting. This work was supported by the Hôpitaux Universitaires de Strasbourg (HUS), by the “Direction de la Recherche Clinique et de l’Innovation” (HUS) and by grants from Strasbourg University (UdS), by the Agence Nationale de la Recherche (ANR-14-CE14-0026-04, Lumugene; ANR-19-CE15-0028, LYMPHO-STING; and ANR-11-EQPX-022). The study was also supported by the Institut National de la Santé et de la Recherche Médicale (INSERM), by the Atip-Avenir, by government grants managed by the Agence National de la Recherche as part of the “Investment for the Future” program (Institut Hospitalo-Universitaire Imagine, grant ANR-10-IAHU-01, Recherche Hospitalo-Universitaire, grant ANR-18-RHUS-0010). Luigi D. Notarangelo is supported by the Division of Intramural Research, National Institute of Allergy and Infectious Diseases, National Institutes of Health.

