## Supplemental Methods and Figures for "Lymphopenia drives T cell exhaustion in immunodeficient STING gain-of-function mice"

**This PDF file includes:**

Supporting materials and methods

Figures S1 to S5

SI references

### SI Materials and Methods

#### Mice and cells

The $STING^{V154M/+}$ mice (called thereafter STING GOF mice) were generated as previously described (1). Mice were bred in specific opportunistic pathogen-free (SOPF) conditions at the animal facility of the Molecular and Cellular Biology Institute (IBMC, Strasbourg, France). Littermate wild-type (WT) and sex-matched mice were always used as control animals. STING GOF mice were crossed with IFNAR1 (IFN-α/β receptor chain 1) knockout (KO) mice, thereafter named IFNAR KO mice (provided by TAAM, CNRS, Orleans, France) (2). All animal experiments were performed with the approval of the " Direction départementale des services vétérinaires " (Strasbourg, France), and protocols were approved by the ethics committee (" Comité Régional d'Éthique en Matière d'Expérimentation Animale de Strasbourg ", CREMEAS) under relevant institutional authorization ("ministère de l'Éducation Nationale, de l'Enseignement Supérieur et de la Recherche "; authorization APAFIS #2387-2015072907553237). The $RAG1^{R972Q/R972Q}$ and $RAG1^{R972W/R972W}$ mice (called thereafter RAG1 hypomorphic mice) have been previously described (3), and experiments with these mice were performed according to protocol ASP LCIM6E, approved by the National Institute of Allergy and Infectious Diseases' animal care and use committee.

Mice were euthanized by cervical dislocation. Spleen, thymus, and bone marrow (BM) were mechanically dilacerated. For pulmonary lymphoid infiltrates analysis, lungs were cut into small pieces and digested with an enzyme solution containing DNase (DN25, 1 mg/mL, Sigma-Aldrich) and collagenase (Liberase, 125 µg/mL, Sigma-Aldrich) for 45 min at 37°C. Red blood cells were removed by osmotic lysis using ACK (Ammonium-Chloride-Potassium) buffer.

#### Flow cytometry

Splenic, thymic, BM or lung single-cell suspensions were analyzed by flow cytometry according to standards protocol. Cells were stained with the following antibodies in PBS (Gibco) supplemented with 2% v/v FBS (Dutscher): anti-CD3e FITC or PE-Cy7 (145-2C11, BD Pharmingen), anti-CD4 AF700 (RM4-5, BD Pharmingen), anti-CD8a FITC, PE or PE-CF594 (53-6 .7, BD Pharmingen), anti-CD44 APC (IM7, BD Pharmingen), anti-CD45.1 APC-Cy7 (A20, Invitrogen), anti-CD45.2 PE-Cy7 (104, eBioscience), anti-CD48 FITC (HM48-1, BioLegend), anti-CD62L PE (MEL-14, BD Pharmingen), anti-CD117 (c-Kit) APC (2B8, BD Pharmingen), anti-CD127 (IL-7Ra) PE-Dazzle594 (A7R34, BioLegend), anti-CD150 (SLAM) PE (TC15-12F12.2, BioLegend), anti-CD223 (LAG3) APC (C9B7W, BD Pharmingen), anti-CD279 (PD1) PE or PE-CF594 (J43, BD Pharmingen), anti-CD366 (TIM3) PE (RMT3-23, BD Pharmingen), anti-Sca-1 (Ly-6A/E) PE-Cy7 (D7, BioLegend), anti-TIGIT APC (1G9, BioLegend).

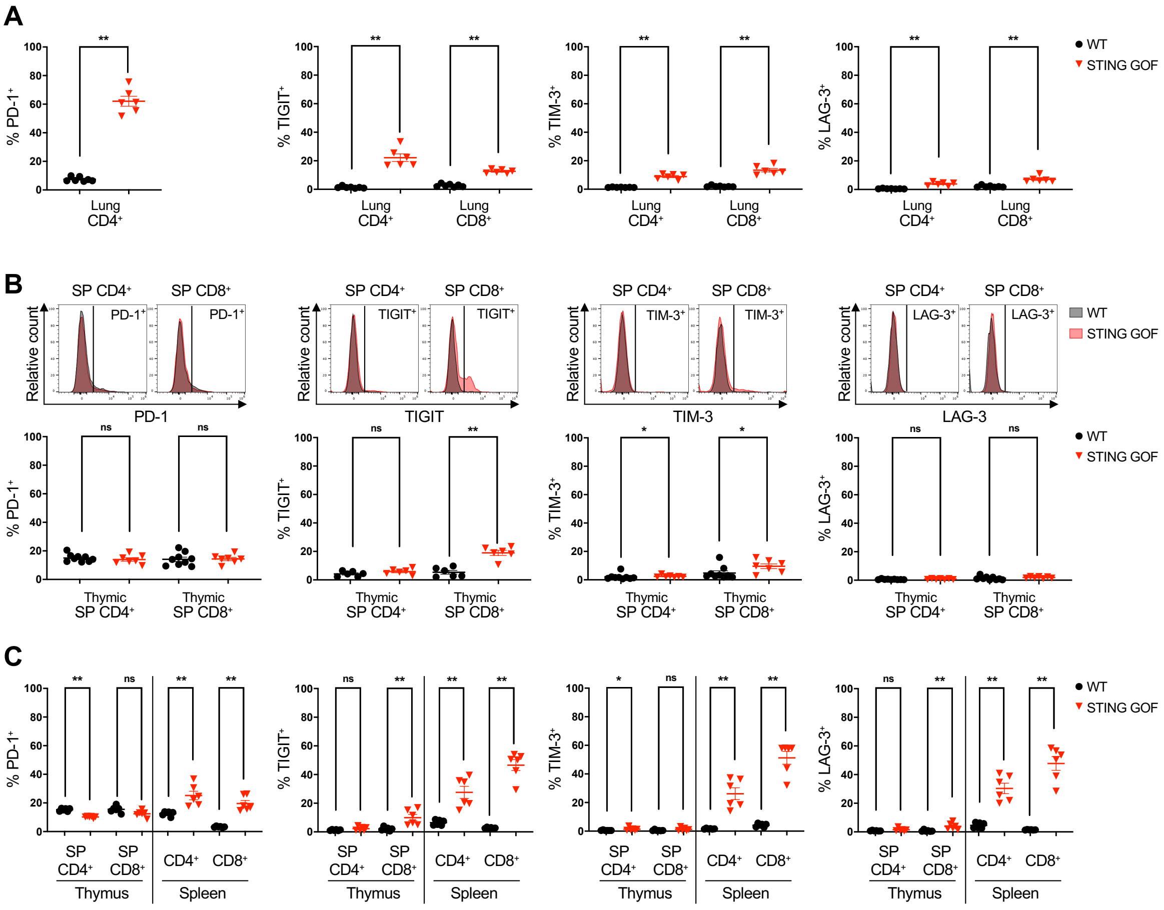


**Fig. S1. T cell exhaustion in STING GOF mice is acquired early in life and in peripheral environment.**

**(A)** Proportion of PD-1-, TIGIT-, TIM-3- and LAG-3-expressing cells among lung $CD4^+$ or $CD8^+$ T cells from STING GOF mice and their WT littermate controls. **(B)** Proportion of PD-1-, TIGIT-, TIM-3- and LAG-3-expressing cells among thymic $CD4^+$ or $CD8^+$ SP T cells from STING GOF mice and their WT littermate controls. Representative histograms are shown. **(C)** Proportion of PD-1-, TIGIT-, TIM-3- and LAG-3-expressing cells among thymic and splenic $CD4^+$ or $CD8^+$ SP T cells from 2-weeks old STING GOF mice and their WT littermate controls. Each data point corresponds to one mouse; mean ± SEM are shown per population for six to nine mice from multiple independent experiments. Statistical significances are calculated with two-tailed Mann-Whitney test: ns, $P > 0.05$; *, $P < 0.05$; **, $P < 0.01$.

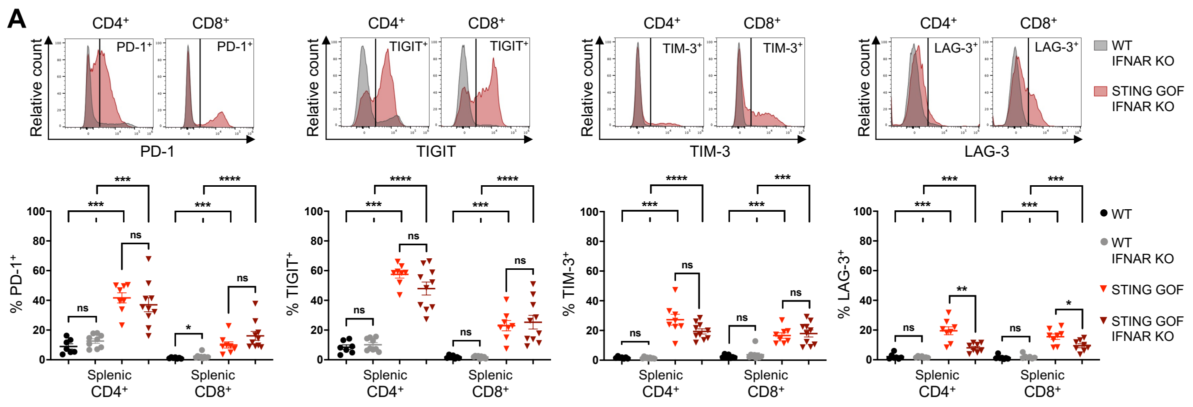


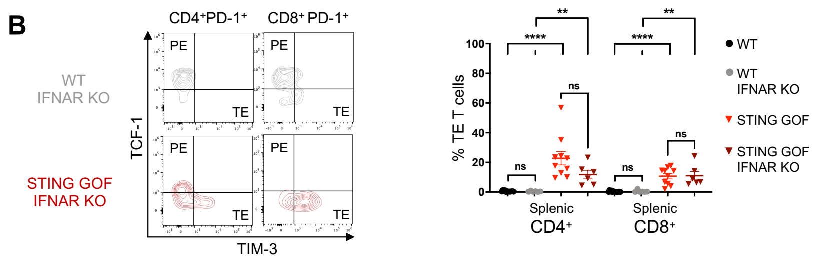


**Fig. S2. T cell exhaustion in STING GOF mice is independent of type I IFNs.**

**(A)** Proportion of PD-1-, TIGIT-, TIM-3- and LAG-3-expressing cells among splenic CD4+ or CD8+ T cells from STING GOF or STING GOF IFNAR KO mice and their respective WT littermate controls. Representative histograms are shown for the IFNAR KO pair. **(B)** Proportion of terminally exhausted (TE) T cells among splenic $CD4^+$ or $CD8^+$ T cells from STING GOF or STING GOF IFNAR KO mice and their respective WT littermate controls. Representative contour plots are shown for the IFNAR KO pair. Each data point corresponds to one mouse; mean ± SEM are shown per population for six to twelve mice from multiple independent experiments. Statistical significances are calculated between IFNAR KO groups as well as between STING GOF groups with two-tailed Mann-Whitney test: ns, $P > 0.05$; **, $P < 0.01$; ***, $P < 0.001$; ****, $P < 0.0001$.

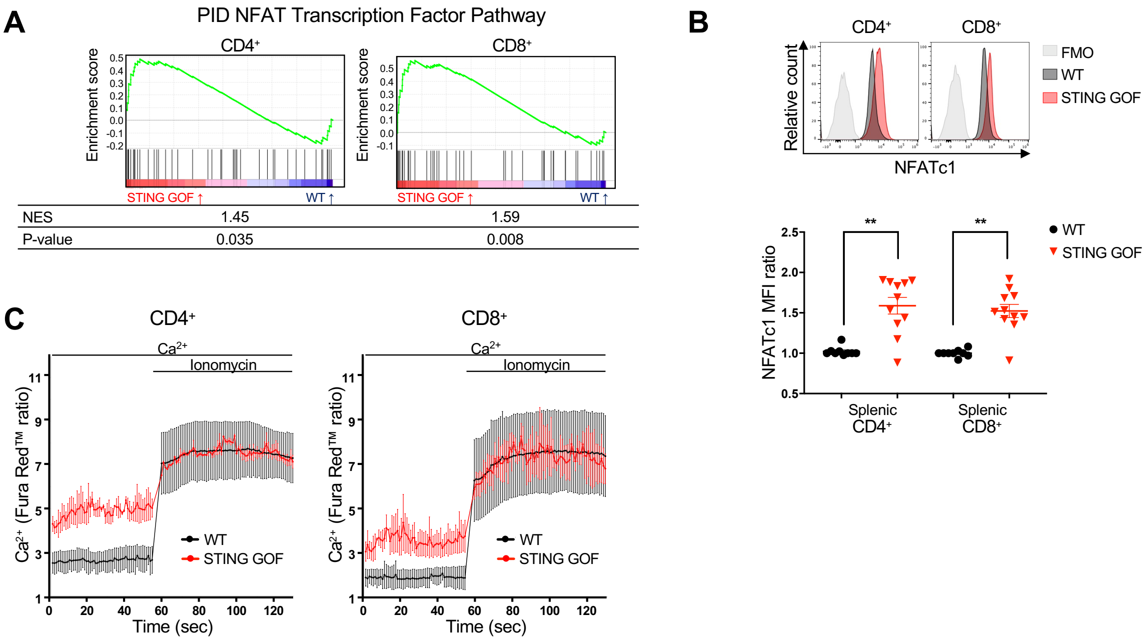


**Fig. S3. $Ca^{2+}$-NFAT activation in STING GOF T cells.**

**(A)** GSEA of the NFAT pathway signature (PID) among genes deregulated in STING GOF versus WT $CD4^{+}$ or $CD8^{+}$ T cells. Enrichment plots, normalized enrichment score (NES) and nominal p-value (p-value) are shown for each analysis. **(B-C)** Immunophenotyping and $Ca^{2+}$levels of splenic T cells from STING GOF mice and their WT littermate controls by flow cytometry. **(B)** Ratio of total NFATc1 mean fluorescence intensity (MFI) in splenic $CD4^{+}$ or $CD8^{+}$ T cells from STING GOF mice and their WT littermate controls. Ratio was normalized on the WT control of each analysis. Representative histograms are shown. Each data point corresponds to one mouse; mean ± SEM are shown per population for nine to eleven mice from multiple independent experiments. Statistical significances are calculated with Wilcoxon signed-rank test with a hypothetical value of 1: **, $P < 0.01$. **(C)** Relative cytosolic $Ca^{2+}$ levels monitored by Fura Red™ ratio in splenic $CD4^{+}$ or $CD8^{+}$ T cells from STING GOF mice and their WT littermate controls. Splenocytes were recorded in DMEM containing 1.8 mM $Ca^{2+}$ and stimulated with 1 µg/mL ionomycin after 50 seconds as positive control. Fura Red™ intensities in VL4 (405 nm-excitation; 660 nm-emission) and BL3 (488 nm-excitation; 695 nm-emission) channels were recorded, corresponding to $Ca^{2+}$-bound and $Ca^{2+}$-free Fura Red™ respectively. Ratio (VL4/BL3) of these Fura Red™ MFI according to time are plotted in a curve graph where each data point represents the mean of two independent experiments, and their +

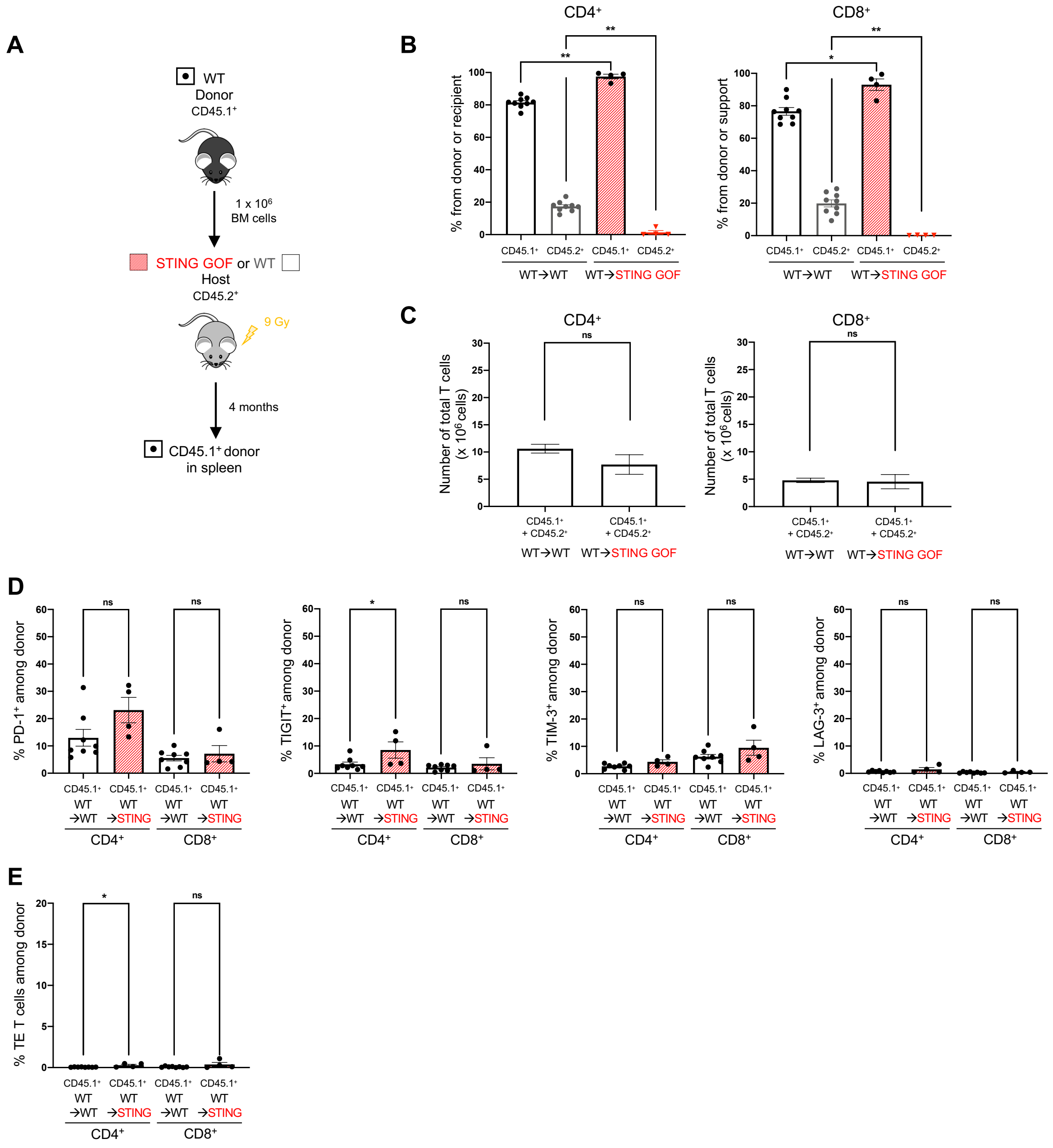


**Fig. S4. STING GOF radioresistant stroma is not sufficient to induce T cell exhaustion.**

**(A-E)** STING GOF mice and their WT littermate controls ($CD45.2^+$) were lethally irradiated and then reconstituted with WT BM donor cells ($CD45.1^+$) to generate WT→STING GOF and WT→WT mice. 4 months later, spleens were assessed for T cell reconstitution and immunophenotyping. **(A)** Strategy of WT BM cells transplantations into STING GOF or WT irradiated recipient mice. **(B)** Proportion of cells derived from WT BM donor cells ($CD45.1^+$) or host cells ($CD45.2^+$) among splenic $CD4^+$ or $CD8^+$ T cells from WT→STING GOF and WT→WT mice. **(C)** Absolute numbers of total ($CD45.1^+$ and $CD45.2^+$) splenic $CD4^+$ or $CD8^+$ T cells from WT→STING GOF and WT→WT mice. **(D)** Proportion of PD-1-, TIGIT-, TIM-3- and LAG-3-expressing cells among splenic $CD4^+$ or $CD8^+$ T cells derived from WT BM donor cells ($CD45.1^+$) from WT→STING GOF and WT→WT mice. **(E)** Proportion of terminally exhausted (TE) T cells among splenic $CD4^+$ or $CD8^+$ T cells derived from WT BM donor cells ($CD45.1^+$) from WT→STING GOF and WT→WT mice. Each data point corresponds to one mouse; mean ± SEM are shown per population for four to eight mice from multiple independent experiments. Statistical significances are calculated with two-tailed Mann-Whitney test: ns, $P > 0.05$; *, $P < 0.05$; **, $P < 0.01$.

A

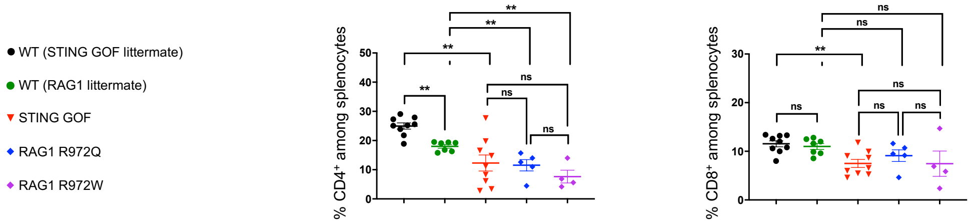


B

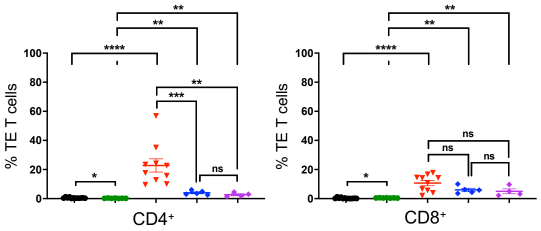


**Fig. S5. Mice carrying *Rag1* hypomorphic mutations also display lymphopenia-associated T cell exhaustion.**

**(A-B)** Immunophenotyping of T cells from the spleen of hypomorphic RAG1 R972Q and R972W mice and their WT littermate controls by flow cytometry. Data were compared with previous result obtained from STING GOF mice and their WT littermate controls (Fig. 1). **(A)** Proportion of splenic CD4$^+$ or CD8$^+$ T cells from hypomorphic RAG1 mice and their WT littermate controls**. (B)** Proportion of terminally exhausted (TE) T cells among splenic CD4$^+$ or CD8$^+$ T cells from hypomorphic RAG1 mice and their WT littermate controls. Each data point corresponds to one mouse; mean ± SEM are shown per population for four to twelve mice from multiple independent experiments. Statistical significances are calculated with two-tailed Mann-Whitney test: ns, P > 0.05; **, P < 0.01; ****, P < 0.0001.
